# A comprehensive manually-curated Compendium of Bovine Transcription Factors

**DOI:** 10.1101/254011

**Authors:** Marcela M de Souza, Juan M Vaquerizas, Adhemar Zerlotini, Ludwig Geistlinger, Benjamín Hernández-Rodríguez, Polyana C Tizioto, Jeremy F Taylor, Marina IP Rocha, Wellison JS Diniz, Luiz L Coutinho, Luciana CA Regitano

**Affiliations:** Federal University of São Carlos, São Carlos SP Brazil; Embrapa Pecuária Sudeste, São Carlos SP Brazil; Max Planck Institute for Molecular Biomedicine, Münster 48149 Germany; Embrapa Informática Agropecuária, Campinas SP Brazil; NGS Genomic Solutions, Piracicaba SP Brazil; Division of Animal Science, University of Missouri, Columbia MO 65211-5300 USA; University of São Paulo, Piracicaba SP Brazil

## Abstract

Transcription factors (TFs) are pivotal regulatory proteins that control gene expression in a context-dependent and tissue-specific manner. In contrast to human, where comprehensive curated TF collections exist, bovine TFs are only rudimentary recorded and characterized. In this article, we present a manually-curated compendium of 865 sequence-specific DNA-binding bovines TFs, which we analyzed for domain family distribution, evolutionary conservation, and tissue-specific expression. In addition, we provide a list of putative transcription cofactors derived from known interactions with the identified TFs. Since there is a general lack of knowledge concerning the regulation of gene expression in cattle, the curated list of TF should provide a basis for an improved comprehension of regulatory mechanisms that are specific to the species.

## INTRODUCTION

Regulation of gene expression is of essential importance for all living species as it controls specific developmental stages and the response to prevailing environmental conditions. The regulation of gene expression also contributes to phenotypic diversity within and between species (1-3).

Among the factors regulating gene expression are proteins known as transcription factors (TFs) that act as initiators of transcription and this class of proteins has been well studied in model organisms. TFs act by recognizing and binding to the regulatory regions of their target genes and can either positively or negatively regulate gene expression (4, 5). TFs bind to specific sequences (motifs) via their DNA-binding domain (DBD) (6). A variety of databases exist that contain collections of protein domain, including Pfam (7), Prosite (8), Smart (9), and Superfamily (10). The InterPro consortium (11) has merged information from these sources and additional 10 databases into entries for protein domains and families. Using the InterProScan tool (12), these domains can be searched for their presence and locations within any assembled genome.

TFs are key proteins in the regulation of important biological processes, for example, embryonic development (13) or tissue differentiation (14). Furthermore, other proteins can interact with TFs to regulate transcription (15). These proteins are called transcription cofactors (TcoFs) and they can form complexes with TFs to fine-tune the precision and complexity of transcriptional regulation.

There have been many studies that investigated human and mouse TFs, their binding domains, target genes, and interactions with other proteins. This has resulted in comprehensive collections of human and mouse TFs (16-23). Among these resources, the human TF census built by Vaquerizas *et al.* (20) includes 1,391 manually-curated human TFs. Additional databases comprise, Animal TFDB (22), DBD (21), and Cis-Bp (23), which provide large collections for 65, 131 and 700 different species, respectively. Animal TFDB also provides a list of TcoFs as derived from known with TFs for each species.

Despite this, knowledge about bovine DNA-protein and protein-protein interactions is limited; TF databases that provide information for the *Bos taurus* exclusively contain TFs that were predicted *in silico* based on data from human and mouse. Although new high-throughput technologies have greatly contributed to a better understanding of gene regulation in cattle, there is currently no curated list of bovine TFs, and all studies in livestock to date have used the human TF list (24-26). Consequently, the development of a compendium of bovine TFs and TcoFs will improve insights into the regulation of gene expression in cattle, reducing opportunities for error caused by humanizing livestock data.

We manually curated a compendium of bovine TFs as derived from the human TF census from Vaquerizas *et al.* (20). We also generated a list of putative TcoFs that have been reported to physically interact with the identified bovine TFs. We are further complementing these collections by analyzing TF evolution, domain family distribution and expression in 14 bovine tissues.

## MATERIALS AND METHODS

### Identification of bovine TF genes

We adapted the approach of Vaquerizas *et al.* (20) by using the property of TFs to bind to DNA in a sequence-specific manner to identify the repertoire of bovine TFs in four main steps (Figure 1).

**Figure 1:**
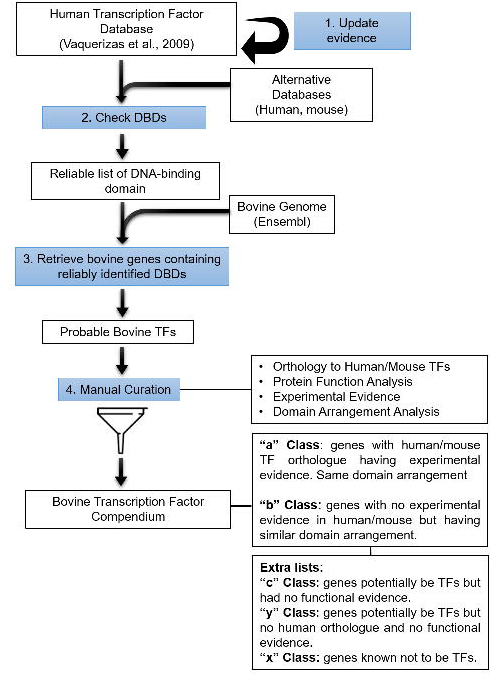
Identification of bovine TFs: 1. Update of the human TF reference repertoire (20); 2. Compilation of reliable DNA-binding domains (DBDs) as in Vaquerizas *et al.* (20), augmented by DBDs found in alternative human and mouse TF databases (AnimalTFDB (22), DBD (21), Cis-BP (23))); 3. Identification of putative bovine TFs using the list of reliable DBDs; 4. Manual curation of the putative bovine TFs by examining orthology to human TFs, protein function, experimental evidence and similarity of domain arrangement. Resulting high-confidence bovine TFs are divided in the evidence classes “a” and “b”.

**Figure 2:**
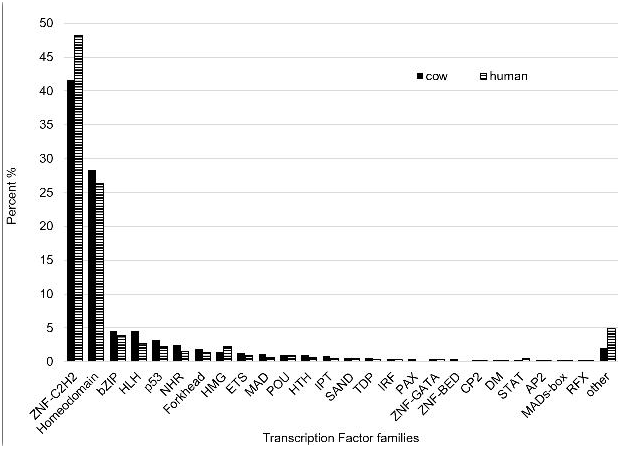
Classification of TFs according to their DNA-binding domain.

*Updating the human reference TF repertoire.* Vaquerizas *et al.* (20) manually curated a list of DNA-binding domains (DBDs), which we updated based on new functional evidence. In the compendium of Vaquerizas *et al.* (20), high-confidence TFs were divided into four classes: “a”-genes that probably encode TFs given experimental evidence for regulatory function in a mammalian organism; “b” - genes that probably encode TFs given an equivalent protein arrangement as for a TF in “a” class; “c” - genes that may potentially encode TFs, but for which there was no functional evidence; and “other” - genes containing unclassified DNA-binding domains obtained from sources such as TRANSFAC (16). Genes known not to be TFs were classified as “x”. Furthermore, we manually inspected TFs initially identified in classes “b” and “c” by Vaquerizas *et al.* (20) to determine if new experimental evidence could be found in the literature allowing their reclassification into “a” class.

*Identification of reliable DBDs.* In the second step (Figure 1), we queried high-confidence TFs (“a” and “b” classes) against three additional human TF databases DBD (21), AnimalTFBD (22) and Cis-Bp (23) to identify probable DBDs that are missing from the Vaquerizas *et al.* (20) list. We first removed from these databases genes classified by Vaquerizas *et al.* (20) as known not to be TFs (“x” class). DBDs common to the three additional databases but that were not contained in Vaquerizas *et al.* (20), were checked for their description and functions reported in the literature. After that, we selected only those domains with a sequence-specific DNA-binding function and that were not found in genes with molecular functions other than transcription. To find new DBDs which may not be present in human, we next applied the same methodology for mouse entries within these three TF databases. We used the InterPro (11) nomenclature for DBDs.

*Identification of probable bovine TFs.* Annotated bovine genes and their InterPro domains from the Ensembl database (release 82) were extracted using BioMart (27). We retained all bovine genes that had at least one DBD contained within the list of reliable DBDs.

*Manual curation.* To remove likely false positives, we compared the predicted bovine TFs with the human counterparts in Vaquerizas *et al.* (20). Ensembl Compara (version 89) was accessed to obtain the human orthologues for all predicted bovine TFs. Bovine genes with one-to-one or one-to-many human TF orthologues of class “a” or “b” in Vaquerizas *et al.* (20) list were selected. We manually inspected all remaining probable TFs by examining the associated literature and selecting those with experimental evidence for either the human or mouse orthologue functioning as a TF. Accordingly selected bovine TFs were classified as “a” class. To ensure that they possessed the same function as their human orthologues (one-to-one or one-to-many) we manually and computationally compared the domain arrangement of each bovine TF against its orthologues, retaining only those with significant domain alignments using the algorithm described by Terrapon *et al.* (28).

Finally, bovine TFs without a human or mouse orthologue were assigned to a new class (“y” class). To check whether those could be bovine-specific TFs, we aligned their DNA sequence to the human genome using Blast (http://www.ensembl.org/Bos_taurus/Tools/Blast). For those showing high sequence similarities to a human gene, we applied a domain arrangement analysis. Resulting bovine genes with a high domain arrangement similarity to a human class “a” or “b” TF were classified as “b”. The remaining predicted bovine TFs without human orthologues were assigned as “y”. Finally, predicted bovine TFs in “y” class included genes without human orthologues but that possessed reliable DBDs. However, no regulatory function was found for them in the literature.

### TF Homology

The evolutionary history of predicted bovine TFs was analyzed using phylogenetic relationships from Ensembl Compara. Orthology information between 21 vertebrates species was accessed using the biomaRt package (29).

### Structural features of TFs

TFs were classified into family groups based on the structure of their DBDs using the same classification scheme as used by Vaquerizas *et al.* (20). TFs with more than one DBD were classified into each of the respective families, and families with less than five members were classified as “other”.

### Identification of bovine TcoF

The TcoF repertoire was built by adapting the approach of Schaefer *et al.* (30). First, proteinprotein interactions were downloaded from IntAct (accessed January 2017) (31). Next, proteins physically interacting with at least one predicted bovine TF were considered as putative TcoFs. Interactions between two TFs were excluded. We filtered for interaction types MI:0195 (covalent binding), MI:0407 (direct interaction) or MI:0915 (physical association).

Putative bovine TcoFs were classified according to their Gene Ontology (GO) annotation. We used the human GO annotation since most bovine annotations are predicted and are not based on experimental evidence from cattle. We required candidate TcoFs to be: i) located in the nucleus (cellular component GO:0005634) and, ii) involved in transcriptional regulation. For the latter, we required molecular functions to include GO:0003713, GO:0003712, GO:0003714, GO:0001221, GO:0001222, GO:0001223, GO:0033613, or GO:0070491 and biological process to include GO:0006351, GO:0045892, GO:0045893, GO:0006355 or GO:0009299. The Entries in the bovine compendium were classified based on GO evidence types. When divided into experimental evidence (EXP, IDA, IMP, IGI, IEP and IPI codes) and non-experimental evidence (all other evidence codes), TcoFs were accordingly classified as “High-confidence” or “Hypothetical,” respectively.

### Tissue-specific expression of bovine TFs and TcoFs

Expression of bovine TFs and TcoFs in 14 tissues was examined using RNA-seq data from the L1 Hereford cow Dominette 01449 described in Whitacre *et al.* (32). Tissues included in the analysis were ampulla, white blood cells, cerebral cortex, endometrium, caruncular regions contralateral (car con) and ipsilateral (car ips) to the corpeus luteum, gallbladder, heart, jejunum, kidney, liver, mesenteric lymph nodes, pons, semitendinosus muscle, and spleen.

Read alignment to UMD3.1 reference assembly was performed using TopHat (33) as described by Tizioto *et al.* (34). In brief, the aligned reads were individually assembled into a parsimonious set of transcripts for each sample. StringTie (35) was used to estimate transcript abundances as Fragments Per Kilobase of exon per Million fragments mapped (FPKM), a procedure that normalizes transcript expression for transcript length and the total number of sequence reads per sample. TF-TcoF co-expression across tissues was analyzed based on simultaneous presence (FPKM > 0) or absence TF-TcoF pairs.

## RESULTS

We curated a compendium of bovine TFs by adapting the approach of Vaquerizas *et al.* (20) in four essential steps (Figure 1).

First, we updated the human TF reference repertoire by inspecting genes classified as “b” or “c” by Vaquerizas *et al.* (20). See Material and Methods for definition of these evidence classes. We found new evidence for transcriptional activity of 86 b-class genes and eight c-class genes, which we accordingly re-classified as “a” (Table S1).

In the second step, we extended the set of high-confidence DBDs from Vaquerizas *et al.* (20) by analysing human and mouse data from three additional TF databases (DB database (21), AnimalTFBD (22) and Cis-Bp (23)). When analyzing human TF data, we found 26 genes that were common to the three additional databases, but that were absent from Vaquerizas *et al.* (20) (Figure S1a). We inspected the DBDs contained in these genes and found zinc finger C2H2-type/integrase DNA-binding domain (IPR013087), which we accordingly added to the set of high-confidence DBDs. When analysing mouse TF data in the three additional databases, we found 1,162 TFs in common (Figure S1b). These TFs contained five novel genes with reliably identified DBDs (IPR001523, IPR008122, IPR008123, IPR017114, and IPR007087) that were also added to the list of high-confidence DBDs. The final list (Table S2) included 133 high-confidence DBDs (corresponding to InterPro entries), which were composed by 76 domains and 57 family domains.

In the third step, we identified probable bovine TFs by searching the collected DBDs in 24,616 bovine genes of the UMD 3.1 genome assembly in Ensembl, extracting 1,525 genes which contained at least one high-confidence DBD.

In the final manual curation step, we obtained human orthologues of the 1,525 predicted bovine TFs from Ensembl Compara. We then removed (i) genes for which human orthologues were classified “c” by Vaquerizas *et al.* (20), (ii) genes not having transcriptional function, and (iii) pseudogenes. This resulted in 1,306 predicted bovine TFs. From these, we further considered putative bovine TFs with orthologues (one-to-one and one-to-many) to human TFs classified as “a”, “b” or “other” by Vaquerizas *et al.* (20). For the remaining genes, for which human orthologues were not present in Vaquerizas *et al.* (20), we analyzed each case for evidence of transcriptional activity in the literature. From this analysis, we recovered four genes that were reclassified as “a” class because we found experimental evidence for TF function for their human or mouse orthologues in the literature. To increase confidence, we verified whether the human orthologues (one-to-one or one-to-many) possessed the same domain arrangement, thereby ensuring that the genes had the same function in the species analyzed. Of the 1,022 predicted bovine TFs analyzed in this step, we found that 865 had identical or highly similar domain arrangements. However, 62 had considerable domain arrangement discrepancies between species. These diverged predicted bovine TFs were excluded from the TF list and classified as “c” along with 95 genes for which we were unable to analyze domain arrangement.

For bovine genes with confidence DBDs but no human orthologues (“y” class), we searched the sequences with BLAST against the human genome assembly GRCh38. We excluded genes with high sequence similarity as well as similar domain arrangement to human genes classified as having functions other than transcription by Vaquerizas *et al.* (20). A total of five genes possessed similarity to genes classified as “a” or “b” by Vaquerizas *et al.* (20) and were classified as “b” in the bovine TF repertoire (Table S3). The remaining 24 genes, without human orthologues and that had reliable DBDs identified but no regulatory function described, were retained in the “y” class (Table S4).

Finally, after analysis of human/mouse orthology, protein function, experimental evidence, sequence similarity, and domain arrangement, the final list of high-confidence bovine TFs contained 865 genes (Table S4 - “a” and “b” classes).

*Comparison to existing bovine TF databases.* We next compared the TFs contained in our bovine TF compendium to those from three existing TF databases (DB database (21), AnimalTFBD (22) and Cis-Bp (23)). This revealed that he majority of TFs in our compendium, 83.2% (N=720), were also annotated as bovine TFs in the three databases. Additional 92 TFs (10.6 %) were present in two, and another 36 (4.2 %) were in only one of the existing databases (Figure S2). Seventeen TFs were exclusively present in our compendium. Of these, 11 were “a” class, with experimental evidence for their TF function, and six were in “b” class. By considering genes that were present within at least one of the alternative sets but that were not in our set, we found evidence for false positive TFs in the above mentioned databases. Of these, 92 had been excluded from our repertoire because they were classified as having other activity than transcriptional by Vaquerizas *et al.* (20) and another 35 in “a” or “b” classes in Vaquerizas *et al.* (20) had domain arrangements that differed from their human/mouse orthologues. Moreover, genes in “a” or “b” classes for which we were unable to perform domain analyses or genes classified as “c” by Vaquerizas *et al.* (20) were included in the alternative TF databases. Genes in these groups require experimental evidence to enable their accurate classification regarding TF functionality.

### TF homology

Using the phylogenetic relationships retrieved from Ensembl Compara, we investigated the presence or absence of orthologues of the 865 predicted bovine TF genes across 21 eukaryotic genomes (Figure 3). We found genes with similar patterns of presence or absence across the species and grouped them in accordance to their conservational similarity. There were 59 (6.8% of the total 865 TFs) TFs that were present only in mammals and another 55 (6.35%) were predominantly found in mammals. Additional 202 (23.35%) TFs predominantly found in vertebrates. From the metazoa TF cluster, (N = 467; 54%), 83 were found in all analyzed species. Finally, 82 (9.5%) TFs were found in most eukaryotes of which, 15 (1.7%) were present in all analyzed species. Interestingly, four TFs were shared by only two species, and 11 TFs had no orthologues in either human or mouse.

**Figure 3:**
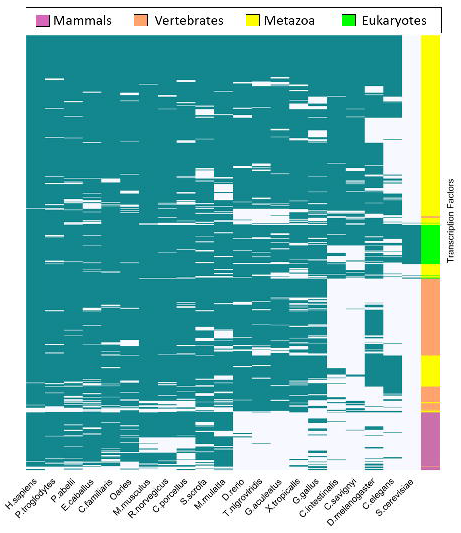
Heat map representation of the conservation of bovine TFs across 21 eukaryotic species. Rows represent the TFs and columns represent the species; both are hierarchically clustered according to the presence (green) or absence (white) of orthologues in the respective species. The color bar on the right indicates whether the TFs are predominantly present in mammals (pink), vertebrates (orange), Metazoans (yellow) or all analyzed eukaryotes (green).

Predicted bovine TFs in the “y” class, which have no human orthologues or evidence of transcriptional function, were also analyzed (Figure S3). We found eight TFs that were present in *Bos taurus* and only one other species, of which two were exclusive to ruminants.

### Structural features of bovine TFs

We grouped the bovine TFs according to their DBD structure, and observed that 76.84% of the TFs belong to four families: C2H2 zinc-finger (n = 596), homeodomain (n = 412), bZip (n = 83) or helix-loop-helix (n = 77). As shown in Figure 2, the distribution of bovine TFs among DBD families was very similar to the distribution of human TF DBD families obtained by Vaquerizas *et al.* (20).

### Identification of bovine TcoFs

We extracted protein interaction data from the IntAct database for all proteins that interacted with the bovine TFs, which resulted in 31,799 interactions. From those, we selected only the 16,608 physical, 1,241 direct and one covalent-binding protein interactions. We inspected each potential TcoF by accessing their GO annotations, to determine if they were located in the nucleus and were annotated to a biological process and molecular function related to transcription. We found 3,842 interacting proteins that were located in the nucleus and 3,590 with GO biological processes, of which 1,558 had GO molecular functions related to transcription. Removing TF-TF interactions yielded 3268 TF-TcoFs interactions of 501 TFs interacting with 782 TcoFs. These TcoFs were classified based on their GO evidence class (Table S5). The highest-confidence class comprised 248 TcoFs with experimental evidence for nuclear localization and molecular function related to transcription. The remaining 534 genes were classified into three groups, called as hypothetical, based on whether they had experimental evidence for nuclear localization, transcriptional function or neither. This resulted in the groups hypothetical I, II and III containing respectvely, 52 proteins with experimental evidence for transcription function but no experimental evidence for nuclear localization, 214 proteins with experimental evidence for nuclear localization but no experimental evidence for transcription function, and 267 proteins with no experimental evidence for nuclear localization or transcriptional function.

### Tissue-specificity of bovine TF and TcoF expression

We next analyzed the expression of the identified TFs and TcoFs as measured with RNA-seq in 14 bovine tissues. Of the 865 TFs and 781 TcoFs in our compendium, 681 (78.7 %) and 608 (77.8 %) were expressed in at least one of the studied tissues, respectively. They were represented by 714 TF (Figure 4A; Table S6) and 635 TcoF isoforms (Figure 4B; Table S7).

**Figure 4:**
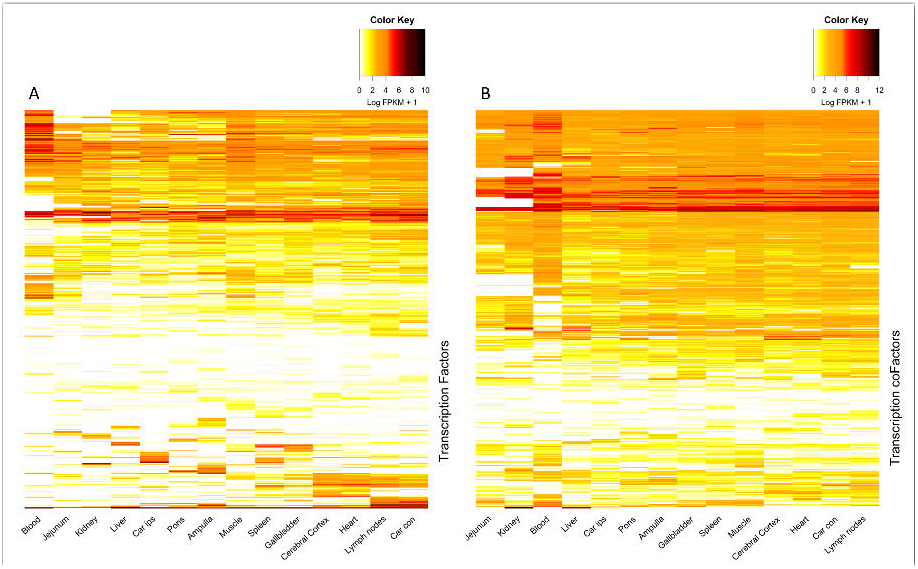
Heat map representation of **(A)** TF and **(B)** TcoF expression in 14 bovine tissues. Columns represent tissues clustered by their expression profile. Each row represents a TF in **(A)** and a TcoF in **(B),** where the color corresponds to the expression level (yellow for low expression, red for high expression, and white for not expressed).

We found considerable variation in TF presence across tissues, ranging from 326 in white blood cells to over 500 TFs expressed in spleen, heart, endometrium sampled from caruncular regions contralateral (car con), lymph nodes, gallbladder, and ampulla. Spleen had the largest number of expressed TFs (N = 541).

Approximately 22.9% of the TFs analyzed were expressed in all 14 tissues, whereas less than 10% were found to be expressed in only one tissue. The Y-box binding protein 1 *(YBX1)* was the most widely expressed TF across all of the tissues, ranging from an FPKM of 18.96 in the ampulla to 882.70 in the kidney. Other TFs expressed in all tissues included *ZFP36* ring finger protein like 1 *(ZFP36L1),* TSC22 domain family member 1 *(TSC22D1),* zinc finger protein 24 *(ZNF24),* X-box binding protein 1 *(XBP1),* DR1 associated protein 1 *(DRAP1),* FOS like 2, AP-1 transcription factor subunit *(FOSL2)* and YY1 transcription factor (*YY1),* which all had an average FPKM of at least 30 across tissues. T-box 20 *(TBX20),* nuclear factor, erythroid 2 *(NFE2)* and T-box, brain 1 *(TBR1)* were exclusively expressed in a single tissue and at high levels (120.83, 58.28 and 22.92 FPKM, in kidney, blood and ampulla respectively).

TcoFs were more broadly expressed than TFs across tissues (Figure S4), with 83.4% of TcoF expressed in more than ten tissues in contrast to only 57.8% of TFs. We also found that 7% of TcoFs but 22.3% of TFs were expressed in at most three tissues. Jejunum had the smallest number of expressed TcoF (N=406) and fewer than 500 TcoFs were expressed in white blood cells and kidney. The other eleven analyzed tissues had between 513 and 567 TcoFs expressed. Heart and spleen (N = 567) had the largest numbers.

We found 40.8% of the TcoFs for which the expression was analyzed to be expressed in all analyzed tissues. The 40S ribosomal protein S3 *(RPS3)* gene was highly expressed in all tissues with an average abundance of expression 617.56 FPKM and ranging from 174.27 FPKM in ampulla to 1,426.03 FPKM in white blood cells. Six other TcoFs were expressed in all tissues and with an average FPKM of 100, and included high mobility group protein B1 *(HMGB1),* 60S ribosomal protein L6 *(RPL6),* prothymosin alpha *(PTMA),* heat shock factor binding protein 1 *(HSBP1),* nucleophosmin *(NPM1)* and elongation factor 1-delta *(EEF1D).* Ankyrin repeat domain-containing protein 1 *(ANKRD1),* and cysteine and glycine-rich protein 3 *(CSRP3)* were expressed in only three tissues but had the greatest expression of all TcoFs (FPKMs of 6,010.14 and 2,863.34 in kidney, 442 and 483.94 in liver, and 2.46 and 0.94 in car con, respectively). Relatively, few TcoFs (2.67%) were exclusively expressed in a single tissue and not at high levels. We found chromobox protein homolog 3 (CBX3) with a FPKM of 19.47 in spleen, and the remaining TcoFs expressed in a single tissue had FPKMs of less than 6.

*TF-TcoF simultaneous expression.* Checking the expression of 2,514 TF-TcoF interaction pairs, we found that 1,937 (77%) TF-TcoF pairs were coexpressed in at least one tissue, from which 278 (11%) were coexpressed in all tissues, and 998 (39.7 %) were coexpressed in more than ten tissues (Figure 5; Table S8). We consider a TF-TcoF pair to be coexpressed when both genes were simultaneously expressed in at least one tissue.

**Figure 5:**
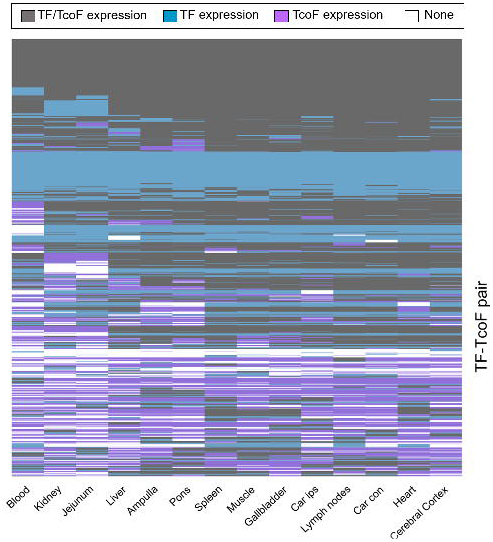
Heat map representation of TF-TcoF coexpression in 14 bovine tissues (white blood cells, kidney, jejunum, liver, ampulla, pons, spleen, semitendinosus muscle, gallbladder, caruncular regions ipsilateral (car ips) to the corpeus luteum, mesenteric lymph nodes, caruncular regions contralateral (car con) to the corpeus luteum, heart, cerebral cortex. Columns represent tissues grouped by their expression profile. Each row represents a TF-TcoF pair.

We found 385 TFs coexpressed with 577 TcoFs. The TF with the most interacting TcoFs, Tumor protein 53 *(TP53),* was coexpressed with 67 of its interacting TcoFs (out of 90, 74.44%). The TcoF with the most interacting TFs, Lysine demethylase 1A *(KDM1A),* was coexpressed with 44 of its interacting TFs (95.65%). The most widely-expressed TcoF, *RPS3* coexpressed with NF-kappaB transcription factor p65 subunit *(RELA)* and *TP53* in all 14 tissues, and with nuclear factor kappa B subunit 1 *(NFKB1)* in 13 tissues.

## DISCUSSION

Knowledge of the existing functional TFs in cattle is of essential importance for studying gene regulatory processes as well as interpreting regulatory implications from high-throughput gene expression data in livestock.

Faced with a lack of information concerning bovine TFs, previous studies (24-26) used the human TF list published by Vaquerizas *et al.* (20) to represent the bovine reference TF set which may lead to errors or oversights. With the availability of a specific bovine TF set, these issues should be minimized and additional insights can be expected in the field of gene regulation in the bovine.

We therefore generated a comprehensive manually-curated compendium of bovine TFs using the human TF census (20) as reference. After updating the human reference, we extended the contained set of DNA-binding domains and searched them in the *Bos taurus* genome sequence assembly. We thereby identified new bovine TFs that were not previously included in existing TF databases.

As existing bovine TF annotation largely relies on orthology transfer from human, it is important to note that we found a non-negligible fraction of human TFs identified by Vaquerizas *et al.* (20) for which the apparent bovine orthologue did not possess the same domain arrangement. As these differences may affect protein function, we excluded putative bovine TFs with predicted domain variation. This also demonstrates that orthology transfer alone is not suficient for accurate bovine TF annotation. For example, the *IKZF2* gene is well described in human and mice as a TF (36) with suggested roles in the regulation of T cell function (36-38) and in the leukemogenesis of adult T-cell leukemia (38). In cattle, we did not find experimental evidences for TF function of *IKZF2* in the literature, and we further found *IKZF2* to have a different domain arrangement than the human orthologue. However, these differences could also partially be artifacts since the bovine assembly is an early-stage draft assembly while the human assembly is essentially complete. Whitacre *et al.* (32) predicted that 42% of bovine genes are either missing or misassembled in the UMD3.1 (32) assembly and this may have produced the domain differences that we found. Thus, we decided to classify these genes as “c” class until further information can be added in the literature and the assembly improved.

Our TF compendium also includes likely bovine TFs without a human or mouse orthologue (and which are thus missing when human TFs are adopted for bovine studies). For example, the gene *LOC509810,* which contains has the same domain arrangement of the human TF *ZNF211* (20), is known to only otherwise be present in sheep and swine. As we also found the gene expressed in 14 bovine tissues, further target studies are needed to clarify the function of this hypothetical TF.

When comparing our results to existing TF databases listing bovine TFs based on orthology transfer from human, the majority of TFs present in our compendium were also included in at least one of the existing TF databases. However, we found that these databases also listed bovine genes as TFs that were excluded from our compendium because of diverged domain arrangements relative to their human orthologues. These databases also listed genes with evidence for functions other than transcription such as *SETDB1* and *SETDB2* that are well-known histone methyltransferases (39, 40). While these genes are classified as TFs in all three the alternative databases, they were classified as not having TF function by Vaquerizas *et al.* (20) and, consequently, were also excluded from our compendium.

Our thorough manual curation of candidate TFs aims at a high-confidence bovine TF compendium. However, it is based on the currently still limited literature for gene regulation in cattle. This is reflected by the incorporated evidence classification scheme. This also allows to distinguish genes containing domains that were confidently predicted as being DNA-binding domains, but that lacked human TF orthologues with an identical domain arrangement. With future studies on gene regulation in cattle, it will presumably become possible to determine if these genes are actually bovine TFs.

To characterize the identified bovine TFs, we checked the presence and distribution of domain families. Although, more recent classification of TF domain families are available in the literature (17), we adopted the same classification scheme as Vaquerizas *et al.* (20) to make a direct comparison possible. As in human (20) and mice (41), the most abundant domain family was C2H2 zinc-finger, followed by homeodomains. This was expected as both domain families are the most common across all eukaryotes, followed by the bZip family (42). C2H2 zinc-finger TFs are only present in eukaryotes (42), whereas homeodomain-containing TFs have also been found in fungi and plants (42).

The evolution of bovine TFs can be assumesd to follow the same pattern as in other mammals (20, 42) as also observed in our results on bovine TF homology to other species.This pattern corroborates the idea that the occurrence of a new type of DBD overlaps with an increment in organismal complexity (43, 44). The notable differences observed for bovine TF orthologues in fungi and the other eukaryotes might can be explained by the evolution of domains such as bHLH (45) and homeodomains (46) after fungi and Metazoa had separated.

The emergence of new domains and their expansions probably enabled an increase in regulatory complexity. For example, Charoensawan *et al.* (42) found that DBD families IRF (interferon regulatory factor) and Churchill (related to neural development) were only present in vertebrates coinciding with the more complex immune and neural systems of vertebrates. Another major expansion occurred with the C2H2 zinc-finger, which is present in both branches of vertebrates and mammals (20, 47). According to Charoensawan *et al.* (42), DBD expansions have been greater in vertebrates than in invertebrates.

We further complemented the compendium by screening for putative transcription co-factors as derived from known interactions with the identified TFs. Using RNA-seq data for 14 tissues from the UMD3.1 reference assembly animal, we analyzed expression profiles for most of the bovine TFs and TcoFs, which suggested that 18% of the TFs and 31.75% of the TcoFs were ubiquitously expressed.

It has previously been shown that genes which evolved early tend to be expressed in more tissues of an organism, whereas more recently evolved genes tend to be tissue-specific in their expression (48). Our results agree with Vaquerizas *et al.* (20) who concluded that TFs do not follow this generalization of an evolutionary pattern of tissue-specific expression. We found TFs that were exclusively expressed in a single tissue but that had orthologues in all analyzed species. Conversely, we found expression of *LOC509810* in all tissues, but this gene apparently has orthologues only in sheep and pig as noted earlier.

Although TF expression analysis was limited to RNA-seq data for a single animal, genes found to be expressed in all analyzed tissues were predominantly housekeeping genes such as *YBX1, ZFP36L1, TSC22D1, DRAP1, FOSL2,* and *YY1.* This is in agreement with results of Harhay *et al.* (49). Despite the majority similarity with the Harhay *et al.* (49) results, *ZNF24* and *XBP1* which we also found to be expressed in all tissues, were not classified as housekeeping by them and, in the opposite, *TBX20, NFE2* and *TBR1* classified as housekeeping genes were expressed in only a single tissue here.

Due to the absence of biological replication and the limited range of tissues represented in the RNA-seq data, general conclusions about the tissue-specificity of TF expression cannot be draw. However, we often found tissue of TF expression to align well with Tf function. For example, *NEF2,* was found to only be expressed in white blood cells in accordance with its function in the maturation of erythroid cells (50, 51) which is delayed when *NEF2* is overexpressed (50). Also, this TF was present in all mammals in our analysis of evolutionary conservation.

In comparison to TFs, TcoFs were apparently more widely-expressed. Around 80% of the TcoFs were expressed in more than ten tissues in contrast to only 57.7% of the TFs. Reciprocally, only 6.9% of the TcoFs were expressed in only one tissue as opposed to 23.7% of the TFs. This can be explained by the fact that each TF may interacts with many TcoFs to initiate transcription and each TcoF can interact with several TFs. Further, we found the TF *TP53* annotated to interact with 90 TcoFs while the TcoF *KDM1A* was annotated to interact with 46 different TFs.

The 40S ribosomal subunit component *RPS3* was the most broadly-expressed TcoF, in agreement with its function as a housekeeping gene (49, 52). We found this TcoF to be coexpressed with three TFs including *RELA* as reported before by Wan *et al.* (52). *RELA,* that was also expressed in all 14 tissues here, is a component of the NF-kB protein complex which act to control the transcription of target genes. The interaction between *RPS3* and *RELA* increases the binding of the transcriptional initiation complex to the DNA (52).

We also analyzed TF-TcoF coexpression to adds experimental evidence to our predictions which were based on GO terms. We found, for example, that the TF *HIF1A,* that is responsive to hypoxia conditions, were expressed in all tissues as so its TcoF *VHL.* In normal conditions of oxigen, *VHL* binds to *HIF1A* preventing the transcription activation of hypoxia-inducible genes (53). However the coactivator *NOTCH1* was not coexpressed with *HIF1A* in any analyzed tissue, which agree with previous studies that the activation of *NOTCH1* transcription is increased in hypoxia condition, and this TcoF directly interact with *HIF1A* in hypoxia-inducible genes promoter (54). However, with no biological replicate we were unable to correlate coexpression profiles whithin each of the tissues for predicted TF-TcoF pairs.

In conclusion, our comprehensive curated bovine TF compendium represents a reliable source of information, with the potential to improve the sensitivity and specificity of studies on gene regulation in the bovine. As we also detailedly characterized the contained TFs with respect to protein structure, evolutionary conservation, and tissue-specific expression, we expect our TF compendium to also be a useful resource for studies on the functions and biological processes in which these TFs are involved. On the other hand, additional experimental evidence for the DNA-binding properties of the TFs in conjunction with additional information about their function and biological activities will also be essential to allow continuous updates and improvements of the compendium.

## SUPPLEMENTARY DATA

### TABLE

**Supplementary Table S1: Updates on evidence for transcriptional activity.** TFs previously classified as “b” or “c” that were reclassified as “a” duo to new evidence of transcriptional activity in the literature. The list contains accompanying information including: Ensembl gene IDs, human orthologue gene, orthology type, bovine TFs repertoire classification, Vaquerizas *et* al. (20) TF classification, literature references of experimentally evidences.

**Supplementary Table S2:** List of Interpro DNA-binding domains and families used to characterise the bovine TFs repertoire.

**Supplementary Table S3: Bovine TFs with BLAST to “a” or “b” class human TF.** The list contains accompanying information including: Ensembl gene IDs, HGNC identifiers, bovine TFs repertoire classification, Vaquerizas *et al.* (20) TF classification, BLAST results.

**Supplementary Table S4: Final list of all genes analyzed and the bovine TFs classification.** List of genes classified as “a”, “b”, “c”, “x”, “y”. The list contains accompanying information including: Ensembl gene IDs, HGNC identifiers, human orthologue gene, orthology type and tissue expression if any.

**Supplementary Table S5: *Bos taurus* TcoF repertoire.** List of TcoF-encoding loci classified as “high-confident” or three “hypothetical” groups. The list contains accompanying information including: Ensembl gene IDs, HGNC identifiers, Uniprot IDs, human orthologue gene, orthology type, bovine transcription factor ID pair, reliability classification and TcoF tissue expression if any.

**Supplementary Table S6:** FPKM *vs* tissue for each TF expressed in at least one tissue.

**Supplementary Table S7:** FPKM *vs* tissue for each TcoF expressed in at least one tissue.

**Supplementary Table S8:** TF-TcoF Co-expression in 14 bovine tissues.

### FIGURE

**Supplementary Figure S1:** Venn diagram comparing TFs from existing transcription factors (TFs) databases. **(a)** Human TFs from Vaquerizas *et al.* (20), Animal TFDB (22), DBD (21) and Cis-BP (23). **(b)** Mouse TFs from Cis-BP, DBD, and TFDB.

**Supplementary Figure S2:** Venn diagram comparing bovine TFs in our compendium with TFs listed for bovine in three existing TF databases: Animal TFDB (22), DBD (21) and Cis-BP (23).

**Supplementary Figure S3:** Heat map showing the conservation of bovine TFs without human orthologues across 20 eukaryotic species. Rows represent the TFs and columns the species; both are hierarchically clustered according to the presence (orange) or absence (white) of orthologues in the respective species. The color bar on the right indicates whether TFs are predominantly present in mammal (pink), vertebrate (orange), Metazoa (yellow) or all analyzed eukaryotes (green).

**Supplementary Figure S4: (A)** Number of TFs and TcoFs, independently determined to be expressed across all tissues. **(B)** Number of tissues in which TFs and TcoFs are independently determined to be expressed.

